# Tau and α-synuclein shape microtubule organization and microtubule-dependent transport in neuronal dendrites

**DOI:** 10.1101/2022.06.09.495530

**Authors:** Marina Rierola, Nataliya I. Trushina, Nanci Monteiro-Abreu, Christian Conze, Michael Holtmannspötter, Rainer Kurre, Max Holzer, Thomas Arendt, Jürgen J. Heinisch, Roland Brandt, Lidia Bakota

**Affiliations:** Osnabrück University, Department of Neurobiology, Osnabrück, Germany; Osnabrück University, Center of Cellular Nanoanalytics, Integrated Bioimaging Facility iBiOs, Osnabrück, Germany; Center for Neuropathology and Brain Research, Paul Flechsig Institute of Brain Research, University of Leipzig, Leipzig, Germany; Osnabrück University, Department of Genetics, Osnabrück, Germany; Osnabrück University, Institute of Cognitive Sciences, Osnabrück, Germany

## Abstract

Tau and α-synuclein are major players in neurodegenerative diseases, but their physiological role, particularly in dendrites, is poorly understood. Here we show that, surprisingly, lack of tau protein induces the development of a more elaborate dendritic arbor of hippocampal pyramidal cells in organotypic tissue. Using high-speed volumetric lattice light-sheet microscopy and single particle tracking, we found a more directional KIF1A-mediated transport in dendrites of Tau KO cells. Increased transport processivity correlated with longer and straighter dendritic microtubules as revealed by three-dimensional super-resolution microscopy of cultured hippocampal neurons. Unbiased mass spectrometric analysis of tissue showed highly increased expression of α-synuclein in Tau KO hippocampi. Overexpression of α-synuclein mimicked the transport characteristics observed in Tau KO cells. Our data indicate that tau and α-synuclein shape microtubule-dependent transport in neuronal dendrites, thereby promoting dendritic arborization during maturation. Furthermore, the data demonstrate that transport efficiency and length and straightness of microtubules are correlated.

## Introduction

Both tau and alpha-synuclein (α-syn) are known for their critical roles in neurodegenerative diseases. Tau is the common denominator in a set of pathological disorders designated as tauopathies. Tauopathies are clinically manifesting with a range of phenotypes that include cognitive, movement, and language disorders and also non-specific amnestic symptoms (Bakota et al., 2017; Zhang et al., 2022). During disease progression, tau is present in a highly phosphorylated state, forming intracellular aggregates that spread through brain regions. The most common tauopathy is Alzheimer’s disease (AD).

With respect to α-syn, point mutations in the α-syn gene (SNCA) or multiplications of the SNCA gene locus lead to the development of monogenetic Parkinson’s disease (PD) (Klein and Schlossmacher, 2006). Neuropathologically, PD is characterized by the progressive loss of dopaminergic neurons of the midbrain in the substantia nigra pars compacta and the deposition of intracellular inclusions in the neuronal cell bodies, referred to as Lewy bodies and Lewy neurites within processes (Goedert et al., 2013). Moreover, concomitant deposition of the two proteins had been extensively described in neurodegenerative comorbidities (Arai et al., 2001; Lippa et al., 1998; Visanji et al., 2019).

α-Syn is a small intracellular protein with a molecular weight of 14 kDa. It has been described as neuron specific in the nervous tissue and is localized in the cytoplasm, presynaptic terminals, and nucleus of the cells (Maroteaux et al., 1988). α-Syn has been shown to be mainly found in the cerebral cortex, cerebellum, striatum, thalamus, olfactory bulb, and hippocampus (Vivacqua et al., 2011).

Tau is a microtubule-associated protein (MAP) that is encoded by the MAPT gene, which in humans comprises 16 exons, located on chromosome 17q21. In the adult brain, tau is expressed in six alternatively spliced isoforms, which are mainly present in neurons, and at lower levels in oligodendrocytes and astrocytes (Kanaan and Grabinski, 2021; LoPresti et al., 1995). Earlier publications suggested that neuronal cells without tau protein would not develop properly, since inhibition of axon formation was observed when using tau antisense oligonucleotides (Caceres and Kosik, 1990). However, other studies using acute tau knockdown in cultured cells or tau knockout mice suggested cooperating and compensatory mechanisms that allow for neuronal cell maturation (Bakota et al., 2017; Dawson et al., 2001; Harada et al., 1994; Tint et al., 1998). There are numerous therapeutical approaches initiated to lower tau protein during neurodegenerative diseases (Dickey et al., 2006; Soliman et al., 2022). However, it is becoming increasingly clear that this can lead to potential side effects (Imbimbo et al., 2022). Therefore, gaining more insights of their physiological role is essential.

Both, tau and α-syn, are mainly suggested and analyzed to play a role in the axon. Tau is enriched in the axonal processes, predominantly at the distal part (Kempf et al., 1996) and α-syn is enriched in the presynaptic terminals (Maroteaux et al., 1988). However, both proteins are also present in the dendrites in smaller amounts (Ittner et al., 2010; Taguchi et al., 2014). The dendritic arbor is a particularly interesting compartment because precise regulation of microtubule dynamics and organization is required to mediate branching that can accommodate the vast majority of synaptic inputs (Penazzi et al., 2016). Dendrites differ from axons in microtubule orientation, stability and polymerization during development of neuronal processes (Kollins et al., 2009). We here set out to understand the dendritic role of tau protein using hippocampal tissue from Tau KO mice by exploring the transport parameters by high-speed volumetric live cell imaging using lattice light-sheet microscopy. Furthermore, we established a three-dimensional super-resolution approach to determine the organization of underlying microtubule tracks within the dendritic processes of cultured hippocampal cells. By utilizing unbiased mass spectrometry, we gained a protein profile comparison of the tissue from Tau KO mice and controls and identified α-syn as a highly overexpressed protein.

## Materials and Methods

### Animals

C57BL/6J mice (onward referred as control, Envigo, The Netherlands) and homozygous TAU^-/-^ mice (Tau KO, (Dawson et al., 2001)) on a C57BL/6J background strain were used for hippocampal primary cultures at embryonic days 16-18, for organotypic hippocampal slices at postnatal day 6-7, and for western blot at 3 weeks of age, and at 3-4 months and at 20-24 months of age. Mice were kept and killed in accordance with FELASA guidelines and approved by the Ethical Committee on Animal Care and Use regulations of Lower Saxony, Germany.

### Human samples

Temporal cortex brain tissue of 11 AD patients (8 female, 3 male) and 10 non-demented age-matched controls (6 female, 4 male) dying without any history of neurological or psychiatric illness was used. Further 6 non-demented cases with an age span 25 to 60 years were included.

Case recruitment, autopsy and data handling have been performed in accordance with the ethical standards as laid down in the 1964 Declaration of Helsinki and its later amendments as well as with the convention of the Council of Europe on Human Rights and Biomedicine and had been approved by the responsible Ethics Committee of Leipzig University (GZ 01GI9999-01GI0299; Approval # 282–02). Informed consent was obtained from all subjects or their legal representatives.

All cases were neuropathologically assessed for neurofibrillary tangle stage according to (Braak and Braak, 1991) and for Aβ/amyloid plaque score according to (Thal et al., 2002) The mean age of the AD cohort (81 +/-6.3 years) was not significantly different from the controls (77 +/-5.4 years). The mean Braak stage of the AD patients was 4.5 +/-0.8 and for age-machted controls 2.1 +/-1.4. The postmortem interval of the AD cohort (65h +/-31h) has been insignificantly longer than that of controls (46 +/-28h) (Table 1).

### Materials

Cell culture media, supplements, culture plasticware, and membrane inserts were obtained from PAN Biotech (Aidenbach, Germany), Thermo-Fisher Scientific (Waltham, USA), Sarstedt (Nümbrecht, Germany), Nerbe Plus (Winsen, Germany), Becton Dickinson Labware (New Jersey, USA), Macherey-Nagel (Düren, Germany), Greiner Bio-One International (Kremsmünster, Austria), Carl Roth (Karlsruhe, Germany), MatTek Corporation (Massachusetts, USA), and Merck KGaA (Darmstadt, Germany), unless otherwise stated.

### Organotypic hippocampal slice cultures

Organotypic hippocampal slice cultures (OHSC) were prepared from 6-7 day-old control or Tau KO mice and processed as described previously (Sundermann et al., 2012; Tackenberg and Brandt, 2009). Briefly, brains were removed from skulls, hippocampi were isolated from the hemispheres and sliced using a tissue McIllwain chopper at 400 µm thickness. Properly shaped slices were selected and transferred onto membrane inserts with fresh culture medium (pH 7.18, Minimum Essential Medium supplemented with 25% HyClone Donor Equine serum, 25% Basal Medium Eagle, 1% glutamine 200 mM, 0.6% glucose, 0.5% Pen-Strep). Tissue was maintained at 37°C and 5% CO_2_. At 11 days *in vitro* (DIV), culture medium was exchanged to supplemented Neurobasal medium (pH 7.18, 1% glutamine 200 mM, 1% N1 supplement, 0.6% glucose, 0.5% HyClone Donor Equine serum, 0.5% Pen-Strep) to ensure viral infection.

### Sindbis virus infection

OHSC were infected using the droplet method (Sundermann et al., 2012) with Sindbis virus to induce the expression of different proteins of interest. To assess morphological parameters, OHSC were infected at 12 DIV and fixed at 15 DIV with ice cold fixing solution (4% paraformaldehyde and 4% sucrose in PBS) and mounted on slides with Confocal Matrix (Micro-Tech-lab, Graz, Austria). To study transport dynamics, OHSC were infected at 13-16 DIV and recordings were made during the following 14-18 hours.

### Sindbis virus preparation

A variety of Sindbis virus constructs based on the triple shuttle vector pJJH1260 were generated by *in vivo* recombination in yeast as described in (Bakota et al., 2012). They were employed for the expression of different genes encoding exogenous proteins within the pSinREP5 vector backbone, as specified in the following. First, freely diffusing enhanced Green Fluorescent Protein (eGFP) was used to assess neuronal morphology in OHSC. In order to analyze transport on OHSC, we used the motor domain (1-396 aa) of the Kinesin Family Member 1A (KIF1A), coding sequence obtained from plasmid Addgene #45058 (Jacobson et al., 2006) C-terminally tagged with mNeonGreen (mNG) and linked via a p2A sequence to either mCherry (pJJH2603), or to mCherry fused to α-synuclein (pJJH2958). Note that 2A self-cleaving peptides are employed to induce ribosomal skipping during translation, enabling the production of two independent peptides from a single mRNA. Finally, a clone encoding a photoactivatable GFP (paGFP) fused to the N-terminus of α-tubulin (pJJH1633) was used to analyze microtubule dynamics in low-density hippocampal primary cultures. Sequences of all viral constructs and detailed strategies of their generation by yeast recombination are available upon request.

### Confocal imaging of neuronal arbors

Tiles of hippocampal neurons localized in the *Cornu Ammonis* 1 (CA1) region were acquired as described previously by (Golovyashkina et al., 2014). Briefly, a Zeiss 510 META NLO confocal laser scanning microscope (Zeiss, Oberkochen, Germany) equipped with a 40x oil-immersion objective (NA, 1.3), a Helium/Neon 543 nm laser and an Argon 488 nm laser was controlled by the LSM 510 v4.0 SP2 software for 3D imaging. Overlapping tiles (7-13 per neuron; voxel size: 0.3 × 0.3 × 0.45 µm) were deconvolved using Huygens Core and the Huygens Remote Manager v3.5 (Scientific Volume Imaging B.V., Hilversum, the Netherlands). Deconvolution was performed using a theoretical point spread function (PSF) and a classical maximum likelihood estimation algorithm with 15 iterations and a quality criterion of 0.05. Overlapping tiles were manually stitched together using VIAS v2.2 software (Mt. Sinai School of Medicine, NY, USA). The neuronal 3D reconstruction of apical and basal arbors was performed by Neuromantic software (University of Reading, Reading, UK) in a semi-automated mode.

### Live-cell imaging of microtubule-based transport mediated by KIF1A under the lattice light-sheet microscope

Lattice light-sheet microscopy was performed on a home-built clone of the original design by the Eric Betzig group (Chen et al., 2014). OHSC were inserted into a custom-built sample holder, which was mounted onto a piezo stage for sample scan imaging. This ensured that the OHSC was inserted at the correct position between the excitation and detection objectives inside the sample bath. The latter contained 7 ml of culture medium at a temperature of 37°C and a saturation of 95% O_2_ and 5% CO_2_. Temperature, gas supply and humidity were precisely controlled by a fully automatic incubation system during the entire experiment (H301-LLSM-SS316 & CO2-O2 Unit BL, Okolab, Italy). Localization of proper CA1 pyramidal neurons in the tissue was granted by the expression of freely diffusing mCherry. In α-syn experiments, mCherry reported proper expression of the protein throughout the neuron. For illumination, a dithered square lattice pattern generated by multiple Bessel beams using an inner and outer numerical aperture of the excitation objective of 0.48 and 0.55, respectively, was used. As illumination sources, a 561 nm laser (2RU-VFL-P-2000-561-B1R; MPB Communications Inc., Pointe-Claire, Canada) was used for mCherry during the identification process, while a 488-nm laser (2RU-VFL-P-300-488-B1R; MPB Communications Inc.) was used during the acquisition of mNG. The final lattice light-sheet was generated by a water dipping excitation objective (54-10-7@488-910 nm, NA 0.66, Special Optics, NJ, USA), while emitted photons were collected by a water dipping detection objective (CFI Apo LWD 25XW, NA 1.1, Nikon, Tokyo, Japan). Emission was then filtered by bandpass filters (Brightline HC 617/73 for mCherry, Brightline HC 525/50 for mNG, Semrock, NY, USA) and imaged on a sCMOS camera (ORCA-Fusion, Hamamatsu, Japan) with a final pixel size of 103.5 nm. For image acquisition, a single-channel image stack was acquired in sample scan mode by scanning the sample through a fixed light sheet with a step size of 400 nm, which is equivalent to a ∼ 217 nm slicing with respect to the z-axis considering the sample scan angle of 32.8°. Imaging was performed in time series of 200 volumes per cell at 600 ms/volume, 850 ms/volume or 900 ms/volume with a frame rate of 100 frames per second based on an exposure time of 8.7 ms per frame plus camera read-out.

### Analysis of dendritic transport mediated by KIF1A

Image series were deskewed, deconvolved, 3D stack rotated, and rescaled with LLSpy (Lattice light-sheet post-processing utility, available at https://github.com/tlambert03/LLSpy). Deconvolution based on the Richardson-Lucy algorithm was performed by using experimental PSFs and 10 iterations. PSFs were recorded from 100 nm sized FluoSpheres™ (Art. No. F8801 and F8803, ThermoFisher Scientific) prior to the experiment. After rotation, images consisted of cubic voxels with a size of 103 × 103 × 103 nm. Files were converted with Imaris File Converter v9.2.1 (Bitplane, Zürich, Switzerland) and kinesin movement marked with mNG was tracked and analyzed using Imaris x64 v9.2.1 (Bitplane, Zürich, Switzerland). First, a reference point, simulating the soma position of the cell, was used to determine the directionality of the trajectories in a dendrite. A ROI containing each dendritic segment was selected and background subtraction was performed. Motor domains were detected by taking into consideration the Gaussian filtered intensity of their signal within a diameter of 300 nm around the signal pick and were tracked semi-automatically in 3D by an autoregressive motion algorithm. This model detects the movement of a trajectory according to the spot detected in the previous volume in order to predict the next spot in the following frame. We set a maximum distance of 3.5 µm, where the next detected spot could be found, and a maximum gap size of 5 volumes. Vesicles detected for less than 3 volumes were discarded. Immobile trajectories (tracks showing a total run length below 0.5 µm) were not further considered in the analysis.

Analysis of directionality and visualization of individual trajectories were performed with R scripts. The total displacement vector of the trajectory showed the general direction of movement. Projections of movement between each tracked point were made onto respective total displacement vectors to determine the mobility of motor domains and the directionality of their movement. The displacement was calculated as a difference between two successive locations of motor domains and normalized by the time between these points. The displacement values lower than 0.2 µm/s were assigned to stationary events. The displacement values higher than 0.2 µm/s were assigned to have “forward” directionality if they aligned with the general direction of the run and “backwards” directionality if they did not align. All changes in movement, for example, stopping or changing directionality, were counted and visualized in displacement plots. The time of motor domains not moving was calculated for all trajectories and averaged by the dendritic segment. The trajectories were considered highly mobile if the total run length was above 15 µm; this allowed the analysis of the processivity of long tracked motor domains and movement changes throughout each trajectory.

### Low-density hippocampal primary cultures

Hippocampi and cortices were separately dissected in ice-cold phenol-red free Hank’s Balanced Salt Solution (Thermo-Fisher Scientific, Waltham, USA) from control and Tau KO mouse embryos at days 16-18 of gestation. Cells were mechanically dissociated and centrifuged at 700 rpm for 15 min at 4°C. To ensure viable isolated neurons, we cultured low-density hippocampal cells and a spatially separated high-dense rim of cortical neurons, as described by (Fath et al., 2009), in previously poly-L-lysine and laminin coated glass coverslip dishes (MatTek, Ashland, MA USA). Supplemented NB/B27 medium was used (Neurobasal medium supplemented with 2% B27, 1% glutamine 200 mM, 1% fetal calf serum, 1% HyClone Donor Equine serum, 0.1% β-mercaptoethanol, 0.2% primocin; or Neurobasal-A medium supplemented with 2% B27, 1% glutamine 200 mM, 0.5% Pen-Strep). Dishes were maintained at 37°C and 5% CO_2_ for 6-8 DIV before usage. In order to measure tubulin dynamics, primary neurons were infected at 5-7 DIV with a Sindbis virus carrying paGFP tagged to tubulin and imaging was performed 20-24 hours post-infection.

### Fluorescent Decay After Photoactivation (FDAP) measurements and analysis

Live imaging was performed using a Nikon Eclipse Ti2-E laser scanning microscope (Nikon, Tokyo, Japan) equipped with LU-N4 laser unit containing 405 nm and 488 nm lasers and a Fluor 60x ultraviolet-corrected oil-immersion objective lens (NA, 1.4) enclosed in an incubation chamber at 37°C and 5% CO_2_. Automated photoactivation was performed with a 405 nm laser in rectangular ROIs of 6 µm length on main and secondary apical dendrites of hippocampal primary neurons at 6-8 DIV from control and Tau KO mice. After activation, a time stack of 1 frame/s was obtained resulting in the acquisition of 112 frames at a resolution of 256 × 256 pixels.

The background subtraction, normalization, and fitting of the FDAP curves were performed with R scripts. For the prediction of the intensity values during photoactivation, the data were modelled using a biphasic exponential decay function:

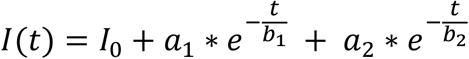

The intensity value directly before image acquisition was calculated using the respective fitting coefficients and substituting the respective time values. Processing and analysis of individual FDAP curves were performed by adapting a reaction-diffusion model from (Igaev et al., 2014). A custom C-based tool called cFDAP was used to model the transformed data and estimate the percentage of polymerized tubulin, and the average association rate k*_on_, and dissociation rate k_off_ of tubulin to and from microtubules. The χ2 value was used as an indicator of the goodness of fit of the model function.

### Sample preparation for DNA-PAINT

Low-density hippocampal cultures at 6-8 DIV were fixed following a NP-40 extraction protocol. Briefly, cells were fixed with a buffer containing 80 mM Pipes KOH, 5 mM EGTA, 1 mM MgCl2, 0.5% (v/v) NP-40, 0.3% (v/v) glutaraldehyde (Sigma G-5882) at pH 8.5 for 10 min at RT. After rinsing with BSA/PBS-Tween20 (1% (w/v) BSA, and 0.1% (v/v) Tween20 in PBS), samples were incubated with 1% (w/v) sodium borohydride (Sigma, S-9125) in PBS for 7 minutes at RT followed by 0.1 M glycine in PBS for 20 minutes at RT. Dishes were rinsed with BSA/PBS-Tween20 and immunostained with monoclonal anti-α-tubulin (1:500, PA5-19489; Invitrogen, MA, USA) o/n at 4°C. Afterwards, cells were washed with PBS followed by the application of an oligonucleotide labeled anti-rabbit nanobody (50 nM, MASSIVE-SDAB 1-PLEX, Massive Photonics, Gräfelfing, Germany) in antibody staining buffer from the same kit at 4°C o/n. Dishes were maintained in PBS at 4°C until imaging was performed. Shortly before image acquisition, cells were incubated with a 1:1000 dilution of 50 nm gold nano rods (E12-50-600-25, Nanopartz Inc., CO, USA) for 5 minutes at RT. Afterwards, samples were washed three times with PBS, after which 3 ml of imaging buffer (Massive Photonics) supplemented with 250 pM of Cy3b labeled imager DNA-strand (MASSIVE-SDAB 1-PLEX, Massive Photonics) were added.

### Three-dimensional DNA-PAINT imaging

For DNA-PAINT, samples were imaged with a total internal reflection fluorescence microscopy (TIRFM) using an inverted microscope (Olympus IX-81) equipped with a motorized quad-line TIR illumination condenser (cellTIRF-4-Line, Olympus, Tokyo, Japan) and a motorized xy-stage (Märzhäuser Scan IM 120×80). Three-dimensional single molecule localization was achieved by astigmatig imaging using a cylindrical lens (Olympus). Imager strands were excited with a 561 nm diode-pumped solid-state laser (max. power 150 mW, Olympus) passing through an 100x oil immersion objective (UAPON 100x TIRF, NA 1.49, Olympus). Excitation intensities were typically adjusted to 20 – 30 W/cm2. Fluorescence emission was filtered by a bandpass filter (BrightLine HC 600/37, Semrock) before being recorded with a sCMOS camera (ORCAFlash 4.0 V3, Hamamatsu). CellSens 2.2 (Olympus) was used as an acquisition software to record 80000 frames with an exposure time of 150 ms per frame and a 2×2 pixel binning which resulted in a pixel size of 130 nm. Sample focus plane was stabilized during acquisition via a hardware autofocus-system (IX2-ZDC2, Olympus). Temperature was kept stable at 25°C with a large incubator chamber (TempController 2000-2, Pecon, Bengal, India) while the sample was humidified via a small stage top incubator (CO2-Controller 2000, Pecon) to exclude buffer evaporation during long-term imaging.

Three-dimensional localization requires axial calibration of astigmatic PSFs by acquiring z-stacks of immobilized fluorescent TetraSpeck™ microspheres with a diameter of 100 nm (T7279; Invitrogen, MA, USA). Stacks were recorded in imaging buffer with a step size of 10 nm using a piezo z-stage (NanoScanZ, NZ100, Prior Scientific, MA, USA).

### DNA-PAINT data analysis

Raw data sets were processed with the “Picasso” software package ((Schnitzbauer et al., 2017), https://github.com/jungmannlab/picasso). First, the calibration z-stack of TetraSpeck™ beads was analyzed with “Picasso localize”. To identify single beads, the box size was set to 13 and the Min. Net. Gradient was adjusted to filter out weak signals from unwanted background localizations. Photon conversion parameters were set as follows: EM Gain: 1, Baseline: 400, Sensitivity 0.46, Quantum Efficiency: 0.80 and pixel size: 130 nm. A calibration file was generated with the “Calibrate 3D” function of “Picasso localize”. This file, as well as the same photon conversion parameters, were used for the image procession of raw sample files. The Min. Net. Gradient was adjusted to remove unspecifically bound imager strands and filter for localizations with the highest signal intensities. Single molecule localizations were fitted with a least square Gaussian fit. For 3D localization, the magnification factor was set to 1.0. Processed datasets were opened with “Picasso render” and a drift correction was first conducted with cross correlation, followed by a correction with gold nano rods as fiducials. The localizations of these data sets where then exported for the ImageJ plugin ThunderSTORM for 3D rendering in order to obtain image-stacks with a well-defined voxel size set to (26 nm x 26 nm x 25 nm) based on the overall axial resolution of 25 nm (Ovesny et al., 2014).

The computational analysis and quantification of the microtubule-array in dendrites of primary neurons was performed by employing the open-source software package SIFNE (SMLM image filament extractor) (Zhang et al., 2017). The MATLAB-based tool involves the iterative extraction of the filamentous structures from the image data set and the subsequent identification and assignment of the detected filaments. Although three-dimensional datasets with a depth between 600 and 800 nm were acquired, we had to create maximum intensity projections (MIPs) from our datasets, as (Zhang et al., 2017) developed their tool for 2D datasets only. Given that the axial spacing between microtubules in dendrites is on average 64±10 nm (Chen et al., 1992) we chose optical sections with a thickness of 100 nm for filament extraction by SIFNE. This allowed us to use sections where the resolution was highest, all while avoiding overlapping superimposed microtubules from another plain that would falsify our statistics. The ROI was set to unbranching dendritic segments of at least 8 µm in length. For image enhancement using line and orientation filter transform (LFT and OFT) algorithms, a radius of 10 pixels with 40 rotations of the scanning line segment was used. Segmentation was performed by employing the automatic threshold functionality of SIFNE based on Otsu’s method. For creating a pool of minimal linear filament fragments, regions of filament junctions were removed by a local region around each junction of 2×2 pixels. In order to recover undetected linear structures, a 2-times iterative extraction of filaments was performed, followed by registration of the propagation direction of each filament tip. Grouping and analysis of detected filaments was performed using a pixel size of 26 nm and a maximum curvature of 1 rad/µm. Search angle and radius were set to 60° and 40 pixels, respectively. Allowable orientation difference between endpoints was set to 60°. The maximum allowable angle difference and endpoint gap vector were set to 60° and 30°, respectively. Weights for similarity and continuity conditions during scoring calculations were set to 1. Due to the high complexity of the cytoskeletal network, no fragment overlap was allowed as suggested by (Zhang et al., 2017) for an intricate network. For sorting of composite filaments, the minimum filament length was set to 15 pixels corresponding to 390 nm, while ungrouped filaments were left in the dataset.

### Sample preparation and peptide generation for Mass Spectrometry (MS)

At 15 DIV of culture maintenance, OHSC were collected, shock frozen and stored at - 80°C until further use. The OHSC were treated with lysis buffer, consisting of 8 M Urea buffer / 50 mM Tris, pH = 7.8 (Sigma-Aldrich Chemie GmbH, Steinheim, Germany; Carl Roth GmbH, Karlsruhe, Germany) and Phos-Stop tablets (Roche Diagnostics GmbH, Mannheim, Germany). Tissue disruption was performed by mechanical pressure as well as tip-sonication for 5 cycles of 5 s each, under 10% amplitude. Next, the samples were centrifuged for 30 min at 4°C and 23,000 x g, pellets containing the organelle debris and insoluble material were discarded and the protein quantification of the supernatant took place using the Pierce™ BCA Protein Assay (Thermo Fisher Scientific, Rockford, USA).

To a protein quantity of 0.03 mg from each sample, 0.2 ug/ul of a-casein solution was added (Sigma-Aldrich Chemie GmbH, Steinheim, Germany). Reduction and alkylation of the proteins was carried out using 8M Urea buffer / 50 mM Tris, pH = 7.8, DL-Dithiothreitol (Sigma-Aldrich Chemie GmbH, Steinheim, Germany) and Iodoacetamide (Sigma-Aldrich Chemie GmbH, Steinheim, Germany), to a final concentration of 5 mM and 15 mM, respectively. Digestion of extracted proteins for peptide generation was performed using Trypsin/Lys-C Mix o/n (Promega Corporation, Madison, USA). The next day, samples were desalted with Pierce™ C18 Tip (Thermo Fisher Scientific, Rockford, USA), followed by lyophilization in a vacuum concentrator. Peptides were dissolved in 0.5% formic acid.

### LC-MS/MS and proteomic analysis

MS analysis was performed as previously described (Schoppe et al., 2020). For full proteome analysis, samples were eluted from a PepMap C18 easy spray column (Thermo Fisher Scientific, Rockford, USA) with a linear gradient of acetonitrile from 10–35% in H_2_O with 0.1% formic acid for 180 min at a constant flow rate of 25 nl/min.

Quantitative proteomic analysis for three samples obtained from Tau KO hippocampi and three samples from control hippocampi was done using PEAKS Online Software (Bioinformatics Solutions Inc., Waterloo, Canada). PEAKS Q (de-novo-assisted Quantification) analysis was used for data refinement with mass correction, de-novo sequencing and de-novo-assisted database search, and subsequent label-free quantification. Mouse proteome from the SwissProt database was used as a target database. Two missed cleavages maximum were allowed per peptide. Carbamidomethylation was set as a fixed modification; oxidation and serine-, threonine-and tyrosine-phosphorylation were set as variable modifications. A maximum of three variable modifications were allowed per peptide. Normalization to the total ion current level for each sample was performed. ANOVA was used to calculate the significance level for each protein, outliers were removed, and the top three peptides were used for protein signal quantification where it was possible.

Proteins identified in only Tau KO hippocampi or control hippocampi were considered uniquely present in the respective condition. Together with significantly up-or down-regulated proteins (p<0.01 and fold change ≥ 2), these proteins were further investigated as differentially regulated proteins.

Gene symbol lists of differentially regulated proteins were uploaded into the Cytoscape software. Significantly enriched GO-terms (p<0.01) associated with biological processes were identified using a right-sided hypergeometric test with Bonferroni correction. GO-term fusion was used to obtain the more representative terms.

### Western blot analysis

Hippocampi from 3 week-old control and Tau KO mice and hemibrains from young (3-4-months-old) and old control and old APP (20-24-months-old) mice were isolated and lysate with ice-cold RIPA buffer (50 mM Tris-HCl, 150 mM NaCl, 1 mM EDTA, 1% NP-40, 0.5% sodium deoxycholate, and 0.1% SDS, pH 8.0) in the presence of protease and phosphatase inhibitors (1 mM PMSF, 10 mg/ml each of leupeptin and pepstatin, 1 mM EGTA, 1 mM sodium orthovanadate, 20 mM sodium fluoride, and 1 mM sodium pyrophosphate). Brain tissue was homogenized in 4 ml/g of the above-mentioned lysing solution. All homogenates were sonicated (10 pulses), and centrifuged for 10 min at 13,000x g at 4°C. Supernatants were frozen and stored at -80°C. Protein concentration of the different homogenates was determined by Pierce_TM_ BCA Protein Assay Kit (Thermo-Fisher Scientific, Waltham, USA). Samples were subjected to SDS-PAGE, transferred to Millipore Immobilon-P PVDF membranes (Thermo-Fisher Scientific, Waltham, USA), and immunoblotting with according antibodies was performed: anti-Tau5 (1:5000, 556319; BD Pharmingen, NJ, USA), α-synuclein (1:1000, PA5-85343; Invitrogen, MA, USA), anti-GAPDH (1:16000, AB2302; Millipore, MA, USA), and, as secondary antibodies, HRP donkey anti-mouse (1:5000, 715-035-151; Jackson ImmunoResearch, Ely, UK), HRP goat anti-rabbit (1:20000, 12-348; Millipore), or HRP donkey anti-chicken (1:10000, 703-035-155; Jackson ImmunoResearch). Protein bands were detected by enhanced chemiluminescence with SuperSignal West Dura (Thermo-Fisher Scientific, Waltham, USA) and quantification was carried out with FusionCapt Advance (Vilber Lourmat, Collégien, France). If required, membranes were afterwards stained during 5 min in 0.1% Coomassie R250 in 50% methanol, 10% acetic acid, followed by a differentiation of the staining in 50% methanol 7% acetic acid to normalize the signal to the total loaded protein amount.

### Immunocytochemistry

Low-density hippocampal cultures at 6-8 DIV were fixed following NP-40 extraction method. First antibodies were applied o/n at 4°C: polyclonal anti-α-synuclein (1:500 PA5-85343; Invitrogen), and monoclonal β-tubulin (1:250 480011; Invitrogen). After rinsing with BSA/PBS-Tween20, secondary antibodies were applied o/n at 4°C: anti-rabbit Alexa Fluor^®^ 488 (1:1000 A11008; Invitrogen), and anti-mouse Cy3 (1:800, 115-165-146; Jackson ImmunoResearch). Dishes were rinsed and imaged in presence of BSA/PBS-Tween20.

### Spinning disk microscopy

To obtain micrographs of immunostained primary hippocampal neurons at 7 and 28 DIV, dual-color high-resolution spinning disk imaging was performed on an inverted microscope (Cell Observer.Z1, Zeiss, Oberkochen, Germany) equipped with a dual camera spinning disk unit (CSU-X1a 5000, Yokogawa, Tokyo, Japan) and a computer-controlled multi-color laser module with AOTF combiner (Zeiss, Oberkochen, Germany). The objective used was an Alpha Plan-Apochromat 63x oil-immersion (NA, 1.46; Zeiss, Oberkochen, Germany), and the acquisition software Zen 2012 (Zeiss, Oberkochen, Germany) was employed. The 488 nm and the 561 nm lasers coupled to an ORCA SNR301410 camera (Hamamatsu Photonics K. K., Hamamatsu, Japan) were used to image Alexa Fluor® 488 and Cy3 fluorophores, respectively. Neurons at 7 DIV were imaged using exposure times of 50 ms (488 nm signal) and 250 ms (561 nm signal). Neurons at 28 DIV were imaged using exposure times of 100 ms (488 nm signal) and 2 s (561 nm signal). 38 HE for green acquisition and a Filterset 43 HE for red acquisition (Zeiss, Oberkochen, Germany) were utilized. Neurons were imaged using a voxel size of 0.086 um x 0.086 × 0.240 µm. All images were deconvolved using Huygens Core and the Huygens Remote Manager v3.5 (Scientific Volume Imaging B.V., Hilversum, The Netherlands).

### Statistical Analysis

Boxplots indicate minimum to maximum data points and the median of the respective condition. All data sets were first checked for outliers, tested for normal distribution using Shapiro-Wilk test, and homogeneity was assessed using Levene’s test. Experiments consisting of two normally distributed conditions were analyzed by independent samples two-tailed t-test; in case variances could not be assumed as equal, Welch’s correction was applied. If normal distribution was not accomplished, Mann-Whitney U test was performed. Transport dynamics mediated by KIF1A experiments involving three conditions were analyzed using one-way ANOVA followed by Dunnett’s post-hoc test if data was normally distributed, otherwise Kruskall-Wallis test was performed. The relationship between variables was assessed with Pearson correlation test. Statistics were performed using SPSS v26 (IBM, NY, USA).

Graphs were generated by GraphPad Prism 8 (GraphPad Software, CA, USA), and R studio (Boston, MA, USA). P values are indicated as follows: * p< 0.05, ** p< 0.01.

## Results

### Tau deficiency increases dendritic complexity in the hippocampus of early postnatal mice

The shape and complexity of dendritic arbor determine the communication in neuronal networks and rely on cytoskeletal modulation. Due to their task of memory formation and retrieval, circuits in the hippocampus are heavily challenged to achieve continuous plastic changes. Therefore, we determined whether the presence of the microtubule-associated protein tau affects the dendritic morphology of hippocampal neurons in cultured tissue environment, which resembles their original connections with other neurons. Organotypic hippocampal slices were maintained for 15 days *in vitro* (Fig. 1A) prior to analysis. The identification, 3D imaging and reconstruction of pyramidal neurons from the CA1 subfield was achieved by virally mediated expression of eGFP in the neurons (Fig. 1B). Analysis of the merged tiles of confocal z-stacks revealed a significantly longer total apical dendritic length of neurons lacking tau (Fig. 1C, left). An increased number of branching points at both apical and basal dendrites was also observed in neurons from Tau KO mice compared to controls (Fig. 1C, right). A more extensive dendritic tree was confirmed by Sholl analysis, which showed more intersections in the middle third of the apical arbor (Fig. 1D). Together, these results indicate an ability to develop increased dendritic branching complexity in the absence of tau protein during neuronal maturation.

**Figure 1.**
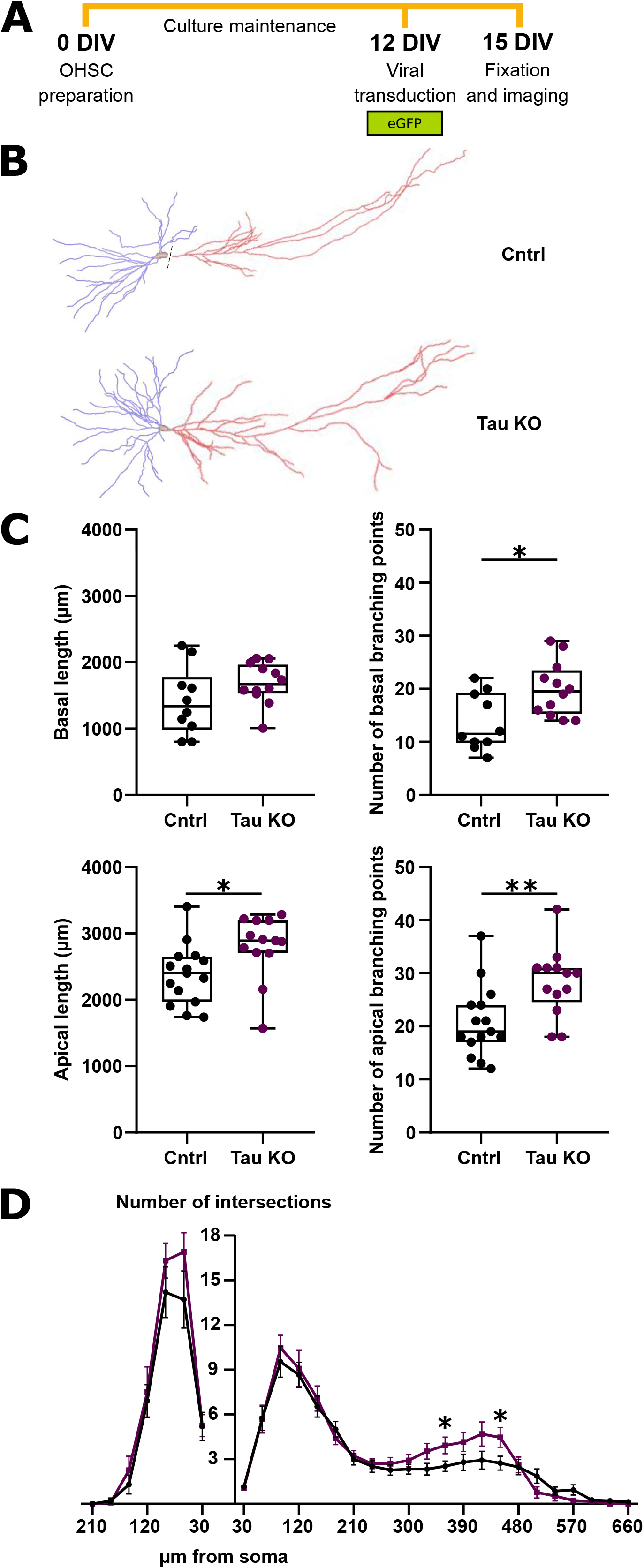
Tau deficiency increases dendritic complexity in the hippocampus of early postnatal mice. **A**. Schematic representation of the Organotypic Hippocampal Slice Culture (OHSC) employed. **B**. Representative reconstruction of basal (blue) and apical (red) dendritic arbors of control and Tau KO pyramidal neurons within the *Cornu Ammonis* 1 (CA1) subregion. Neurons expressed enhanced Green Fluorescent Protein (eGFP) through a Sindbis virus vector for visualization. **C**. Quantification of arbor length and number of branching points at the basal and apical trees. **D**. Sholl analysis of basal and apical dendritic arbors indicating number of intersections along equal sections of µms from the soma. n = 10 control basal arbors; n = 12 Tau KO basal arbors; n = 15 control apical arbors; n = 13 Tau KO apical arbors. Statistical analysis was determined by unpaired two-tailed t-test. *p<0.05; **p<0.01.

### The dynamics of the dendritic microtubules are not affected by the lack of tau protein

Tau is a microtubule-associated protein that regulates the tubulin-microtubule balance in neurons. Therefore, we analyzed whether the lack of tau protein would affect the dynamics of dendritic microtubules, which could influence arbor formation. To determine possible changes in microtubule dynamics, we used a previously established photoactivation assay (Conze et al., 2022). Viral mediated expression of photoactivatable GFP-tagged α-tubulin was performed in primary hippocampal cell cultures prepared from control or Tau KO mice (Fig. 2A). A population of the tagged tubulin was focally activated in the dendrites of pyramidal shaped neurons and the fluorescence decay after photoactivation (FDAP) was monitored over time in the region of activation (Figs. 2A and B). FDAP curves were very similar in control or Tau KO neurons (Fig. 2C, left). To determine the fraction of free and bound tubulin, we fitted the FDAP curves with a reaction-diffusion model. The amount of polymerized tubulin was ∼70% and did not differ significantly between control and Tau KO cultures (Fig. 2C, right). Application of the reaction-diffusion model revealed k*_on_ and k_off_ values for average microtubule polymerization that were slightly increased in cells from Tau KO mice (Fig. 2C, middle), indicating a small increase in microtubule dynamics when tau is lacking. However, the changes did not reach a significance level. Thus, the data show that the absence of tau does not affect the dynamics of dendritic microtubules to a major extent.

**Figure 2.**
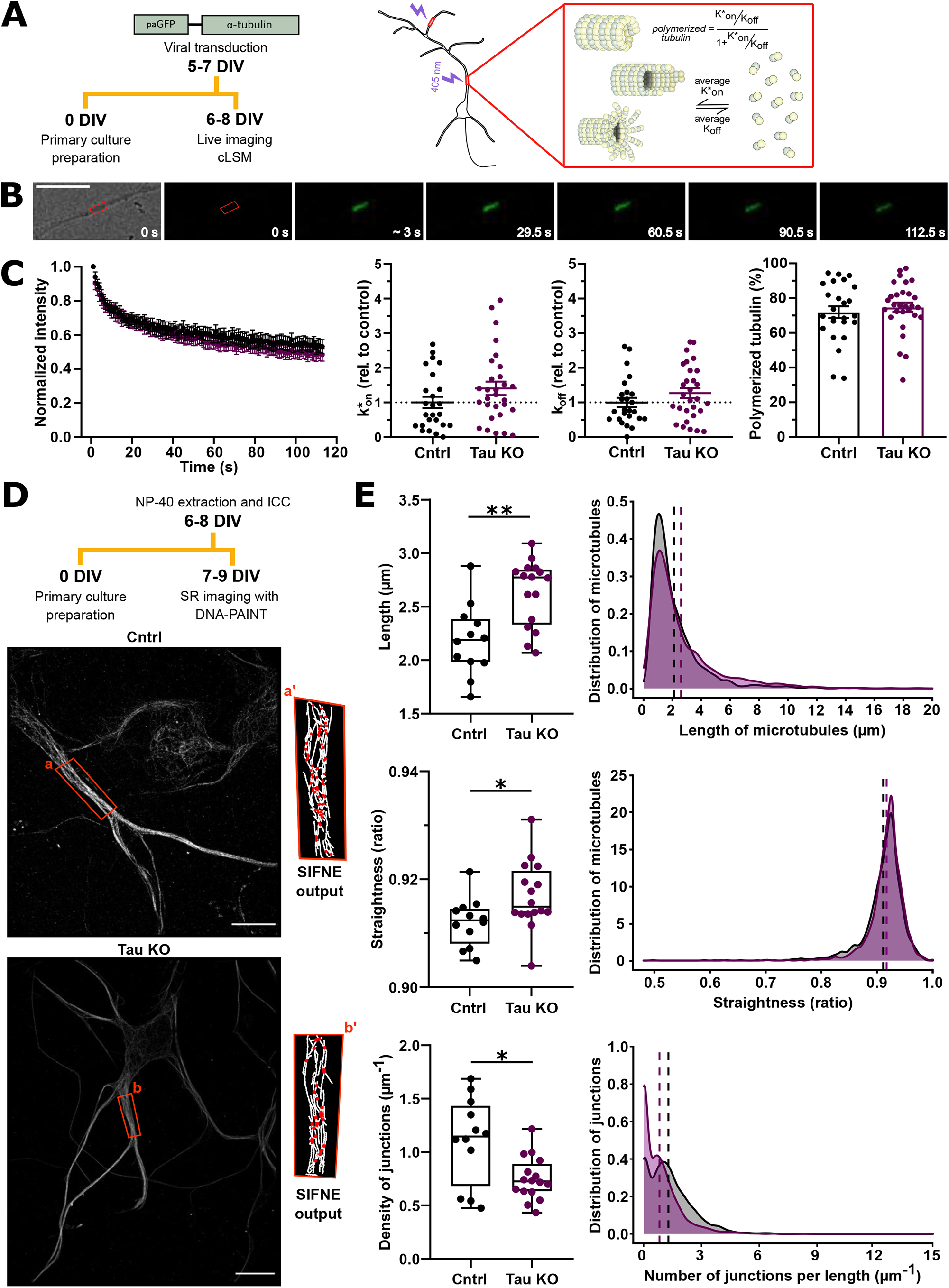
The dynamics of the dendritic microtubules are not affected by the lack of tau protein but the structure of microtubules is altered. **A**. Schematic representation of the method employed (left; plasmid used: pJJH1633). Pictogram of a hippocampal primary neuron, where a segment of the main dendrite and a segment of a secondary dendrite are shown to be photoactivated by a 405 nm wavelength laser (middle). Conceptual representation of the estimated polymerization of tubulin, and association (k*_on_) and dissociation (k_off_) rates determined by Fluorescence Decay After Photoactivation (FDAP) experiments (right). **B**. Exemplary time-lapse images of a dendrite, where the exogenously expressed paGFP-α-tubulin was photoactivated in a 6 µm long ROI (red box). The fluorescence decay within the region, was monitored at a speed of 1 second/frame. Scale bar: 20 µm. **C**. Decay curves displaying mean values of normalized intensity measurements (left). Estimated average of association (k*_on_) and dissociation (k_off_) rates (middle) and bar plot showing the estimated percentage of polymerized tubulin (right). Data are displayed as mean ± SEM. n = 25 control and n = 29 Tau KO dendritic segments from hippocampal primary neurons. Statistical analysis employed unpaired two-tailed t-test. No significant differences were encountered. **D**. Schematic representation of the method used to perform SR (super-resolution) imaging with DNA-PAINT (DNA points accumulation for imaging in nanoscale topography). Below, exemplary control and Tau KO neurons imaged with DNA-PAINT targeting α-tubulin. Selected ROIs (a, b) were processed with SIFNE (single-molecule localization microscopy image filament network extractor) to obtain information about individual microtubules and junctions (red dots in a’ and b’). Scale bar: 10 µm. **E**. Box plots indicating total length of microtubules, straightness, and density of junctions in control and Tau KO dendrites (left). Distribution curves of the individual microtubules according to their length, straightness and number of junctions (right). n = 12 control and n = 16 Tau KO hippocampal primary neurons analyzed with 1400 and 1485 microtubules, respectively. Statistical analysis employed unpaired two-tailed t-test. *p<0.05; **p<0.01.

### Microtubules reorganize into longer and straighter tubules in dendrites lacking tau protein

Although tau has not significantly altered the dynamics of dendritic microtubules, it can still affect their structural organization. To study this, we used a super-resolution imaging approach to quantitatively assess the structure of the microtubules in the dendrites. We employed 3D DNA-PAINT (DNA-based point accumulation for imaging in nanoscale topography), which exploits transient hybridization between oligonucleotide strands and enables single molecule localization with a lateral resolution below 10 nm, and an axial resolution below 50 nm, respectively (Jungmann et al., 2010). Microtubules in the dendritic processes of cultured hippocampal cells were tagged by immunolabeling using a secondary antibody conjugated to a specific single-strand DNA “docking” oligo. Using TIRF microscopy in the presence of low nanomolar concentration of labeled “imager” oligos (Cy3B) allowed for three-dimensional single molecule localization microscopy (SMLM) of microtubule filaments with nanometer resolution (Fig. 2D). Rendered super-resolved z-stacks were further analyzed using the open-source software package SIFNE (single-molecule localization microscopy image filament network extractor) (Zhang et al., 2017). Microtubules in cells from Tau KO mice showed a significant increase in length compared to microtubules in dendrites of control cells (Fig. 2E, top left). The increase was due to a population shift of microtubules towards longer ones (Fig. 2E, top right). The microtubules were also straighter (Fig. 2E, middle) and less intertwined, indicated by a lower density of junctions representing a crossed array of adjacent microtubules (Fig. 2E, bottom). Thus, the data show that microtubules in the dendrites of hippocampal neurons are longer, straighter, and more parallel in the absence of tau.

### Lack of tau protein initiates strong overexpression of α-syn

To gain a broader understanding of the changes that might contribute to the differences in microtubule organization between the two genotypes, we took an unbiased approach by applying mass spectrometry to compare protein profiles of cultured hippocampal slices. Functional analysis of differentially regulated proteins revealed changes in Tau KO animals associated with neuron development, exocytosis, regulation of apoptosis, post-translational regulation, and cytoskeleton organization (Supplementary Fig. 1). We did not observe compensatory overexpression of other microtubule-associated structural proteins (Fig. 3A, top left). However, we found that some motor proteins of both the kinesin and dynein families were upregulated in the tissue sections from Tau KO mice (Fig. 3A, top right). By far the greatest change was a massively increased expression of α-synuclein (α-syn) in Tau KO samples compared to controls (Fig. 3A, bottom). By western blot analysis, we confirmed a strong ∼5-fold overexpression of α-syn in 3-week-old hippocampi (corresponding to the age of the cultured slices) of Tau KO mice (Fig. 3B).

**Figure 3.**
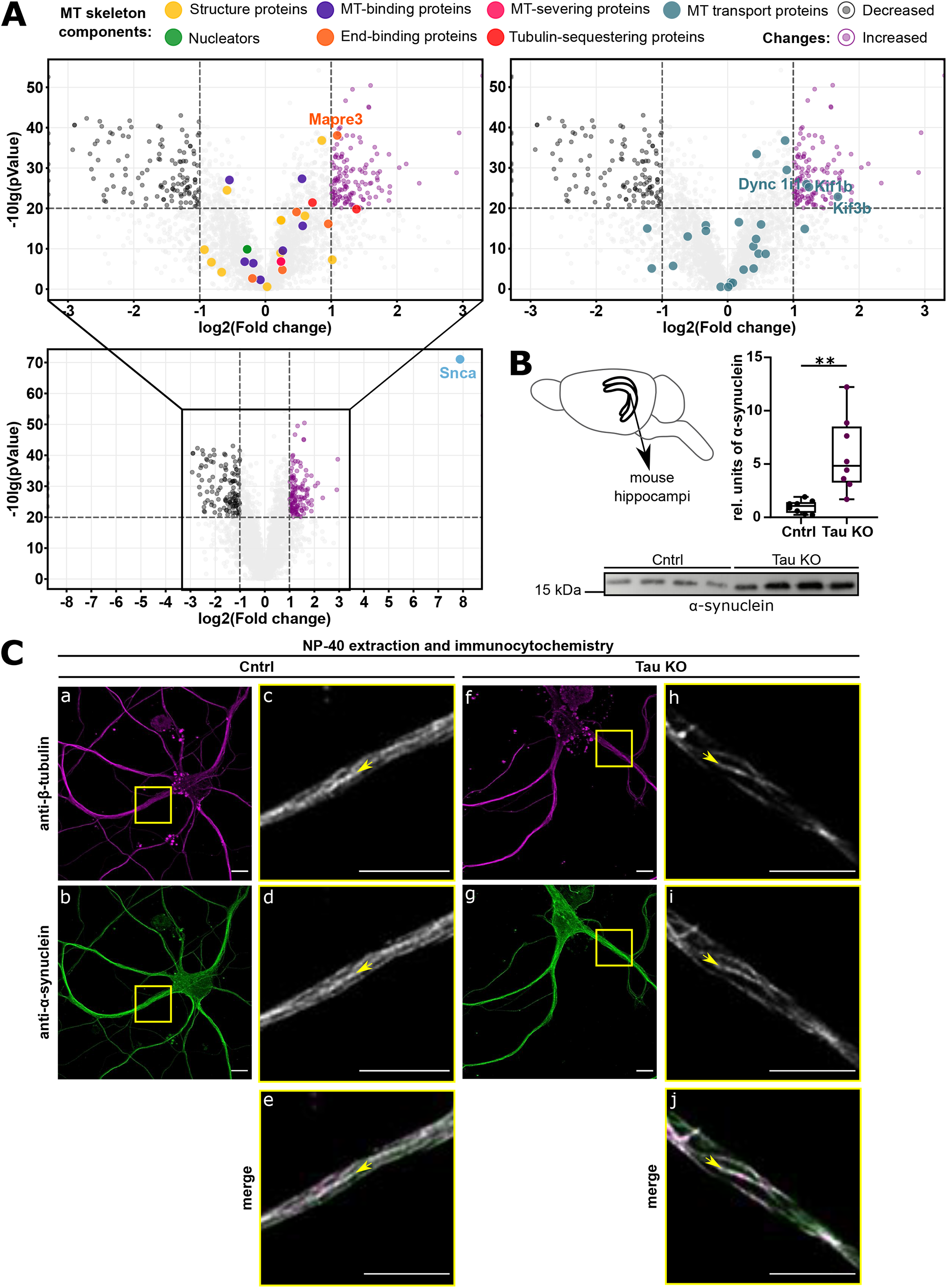
α-syn expression is up-regulated in the absence of tau. **A**. Volcano plots indicating proteins detected by mass spectrometry in OHSC from Tau KO mice in comparison to control. Up-and down-regulated microtubule structure modifying proteins (left) or microtubule transport proteins (right) are highlighted. Below, the volcano plot axes were enlarged in order to reveal that α-syn expression is highly increased in the absence of tau. **B**. Pictogram representing the localization of hippocampi in mouse brains. Box plot indicating the relative amount of α-syn measured and normalized to the protein load in control and Tau KO hippocampal lysates (right). Below, representative western blot bands showing α-syn expression in control and Tau KO lysates. The molecular mass standard is indicated. n = 8 control and n = 8 Tau KO hippocampal lysates from 3-week-old mice. **C**. Exemplary micrographs showing the z-projection of a hippocampal primary neuron immunostained with anti-β-tubulin and anti-α-syn antibodies, from control and Tau KO mice (a,b,f,g). A z-plane displaying the inner segment of the dendrite was selected (c,d,h,i). Below, selected z-planes where merged to show co-localization of β-tubulin and α-syn signal (e,j). Yellow arrows indicate microtubule bundles where co-localization can be observed. Scale bar: 10 µm.

*In vitro* studies have previously shown an interaction between tubulin and α-syn (Alim et al., 2002; Cartelli et al., 2016) that may contribute to the alteration in microtubule organization that we observed in neurons lacking tau protein. To determine a possible interaction of α-syn with dendritic microtubules, we performed a combined NP-40 permeabilization-fixation protocol that preserves the association of MAPs with microtubules (Lee and Rook, 1992), followed by double staining of microtubules and α-syn. Anti-tubulin staining showed the microtubule network in cultured hippocampal cells from wild-type and Tau KO mice. Staining against α-syn revealed the well-described presynaptic accumulation characteristic of its distribution in excitatory neurons (Supplementary Fig. 2). However, we also found that α-syn clearly decorated microtubule bundles in the dendritic processes of cells of both genotypes (Fig. 3C). Therefore, our data suggest that α-syn contributes to the organization of dendritic microtubules and may also modulate microtubule-dependent functions, particularly in a tau protein-free scenario.

### Lack of tau increases the efficiency of dendritic transport

Reorganization of dendritic microtubules can facilitate cargo transport, resulting in the more elaborate dendritic arbor seen on CA1 neurons from Tau KO mice. To test this hypothesis, we examined motor protein dependent cargo transport in dendrites of CA1 pyramidal neurons in tissue culture using high-speed volumetric lattice light-sheet microscopy. We prepared a viral construct encoding the KIF1A motor domain labeled with mNeonGreen (mNG) (Shaner et al., 2013) followed by a self-cleaving peptide sequence (p2A; (Liu et al., 2017)) and mCherry (Fig.4A, left). After cell transduction, the freely diffusing mCherry allowed identification of the respective neuronal cells within the hippocampal tissue (Fig. 4A, right). The bright monomeric green fluorescent protein, mNG, enabled the movement of individual motor protein-driven vesicles to be tracked within the dendrites (Fig. 4B). To quantify vesicle transport, we determined velocity, processivity and speed based on the individual vesicle trajectories analyzed from each cell. Transport via KIF1A showed significantly increased vesicle processivity in neurons lacking tau protein without a change in velocity and speed of vesicles in the apical dendrites of CA1 neurons from Tau KO mice compared to controls (Fig. 4C). Analysis of the individual trajectories revealed no difference in the pause time, but significantly fewer directional changes in the dendrites of cells without tau (Fig. 4D). The results suggest that lack of tau increases the efficiency of dendritic transport via KIF1A, which could facilitate dendritic arborization in the CA1 neurons of Tau KO mice.

**Figure 4.**
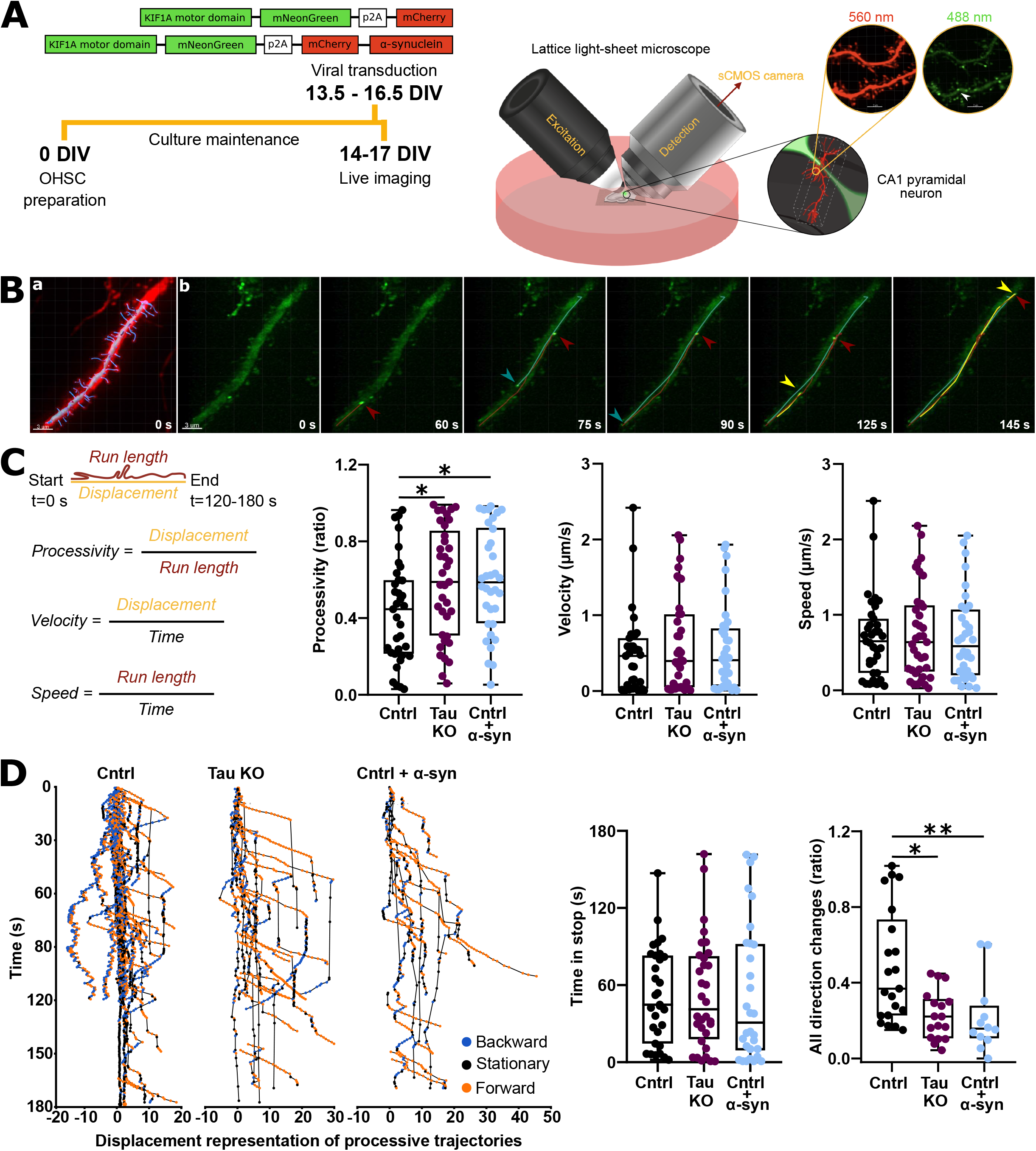
Lack of tau increases the efficiency of dendritic transport. **A**. Schematic representation of the OHSC method employed (plasmids used: pJJH2603 and pJJH2971). Pictogram representing the core of our home-built lattice light-sheet microscope (LLSM) and representation of a localized CA1 pyramidal neuron in the tissue imaged. The 560 nm and the 488 nm wavelength boxes display examples of neuronal dendrites imaged. The white arrow head placed within the 488 nm wavelength box indicates an example of a trackable vesicle. Scale bar: 2 µm. **B**. Time-lapse micrographs of a representative CA1 pyramidal neuronal dendrite imaged with LLSM. Expression employed a Sindbis virus vector coding for the motor domain of kinesin 1A (KIF1A) tagged to mNeonGreen (mNG) followed by a self-cleavage p2A sequence and mCherry. Freely diffusing fluorophore mCherry (a - displayed in red) was used in order to localize CA1 neurons in OHSC and visualize the morphology of dendrites of interest (a - dendritic shaft and dendritic spines reconstructed in grey and blue). When α-syn was overexpressed in control OHSC, mCherry visualization was also used as a reporter. The motor domain of KIF1A tagged to the mNG fluorophore allowed to track vesicle trajectories in dendrites (b). Indicated in burgundy and yellow, examples of vesicles showing transport towards the dendrite terminal are highlighted. In blue, a transported vesicle towards the cell body is displayed. Scale bar: 3 µm. **C**. Quantification of processivity (ratio of displacement and run length for a measured trajectory), velocity displacement of a trajectory per time), and speed (run length of a trajectory per time) per dendrite. n = 35 control, n = 37 Tau KO, and n = 34 control overexpressing α-syn dendritic segments containing 143, 136, and 95 trajectories, respectively. Statistical analysis of processivity was determined by One-way ANOVA followed by Dunnett’s post-hoc test, and velocity and speed statistical analysis were performed by non-parametric Kruskal-Wallis test. *p<0.05. **D**. Individual highly processive trajectories are represented according to time and relative displacement in order to observe differences in their mobility (left). Box plot displaying the amount of time vesicles remained stopped in highly mobile trajectories analyzed. n = 31 control, n = 32 Tau KO, and n = 30 control overexpressing α-syn dendritic segments containing 140, 132 and 75 trajectories, respectively. Box plot displaying the ratio of direction changes. n = 21 control, n = 17 Tau KO, and n = 12 control overexpressing α-syn dendritic segments containing 105, 79 and 45 trajectories, respectively Statistical analysis was performed by non-parametric Kruskal-Wallis test. *p<0.05; **p<0.01.

### Elevated levels of α-syn are sufficient to increase the efficiency of dendritic transport in control cells

Since tau deficiency was associated with elevated α-syn, we next analyzed the effect of increased expression of α-syn in control mice. To achieve this, we prepared an additional viral construct encoding the mNG-labeled KIF1A motor domain followed by mCherry-tagged α-syn after the self-cleaving peptide sequence (Fig.4A, left). After cell transduction, velocity, processivity and speed of vesicle transport were analyzed as previously described. Similar to the experiments with Tau KO cells, we observed no difference in velocity and speed but a significant increase in the processivity of transport (Fig. 4C). Here, too, the analysis of the individual trajectories showed no difference in the pause time, but fewer changes in direction (Fig. 4D). Thus, the data suggest that the elevated α-syn levels are responsible for the increased efficiency of dendritic transport observed in neurons from Tau KO mice.

### Tau and α-syn levels show a negative correlation in aged mice

The high overexpression of α-syn in mice lacking tau protein could indicate a negative correlation between tau and α-syn levels even under physiological conditions. To test this hypothesis, we prepared lysates from hemibrains of young and old control mice and subjected them to western blot analysis (Fig. 5 A). Tau levels in young mice were restricted to a narrow range and we observed a positive correlation between α-syn and tau levels at this age stage. In old mice, tau levels were significantly reduced, highly variable, and we observed a clear negative correlation between tau and α-syn. Thus, the data may indicate that decreased tau levels during aging are compensated by increased expression of α-syn.

**Figure 5.**
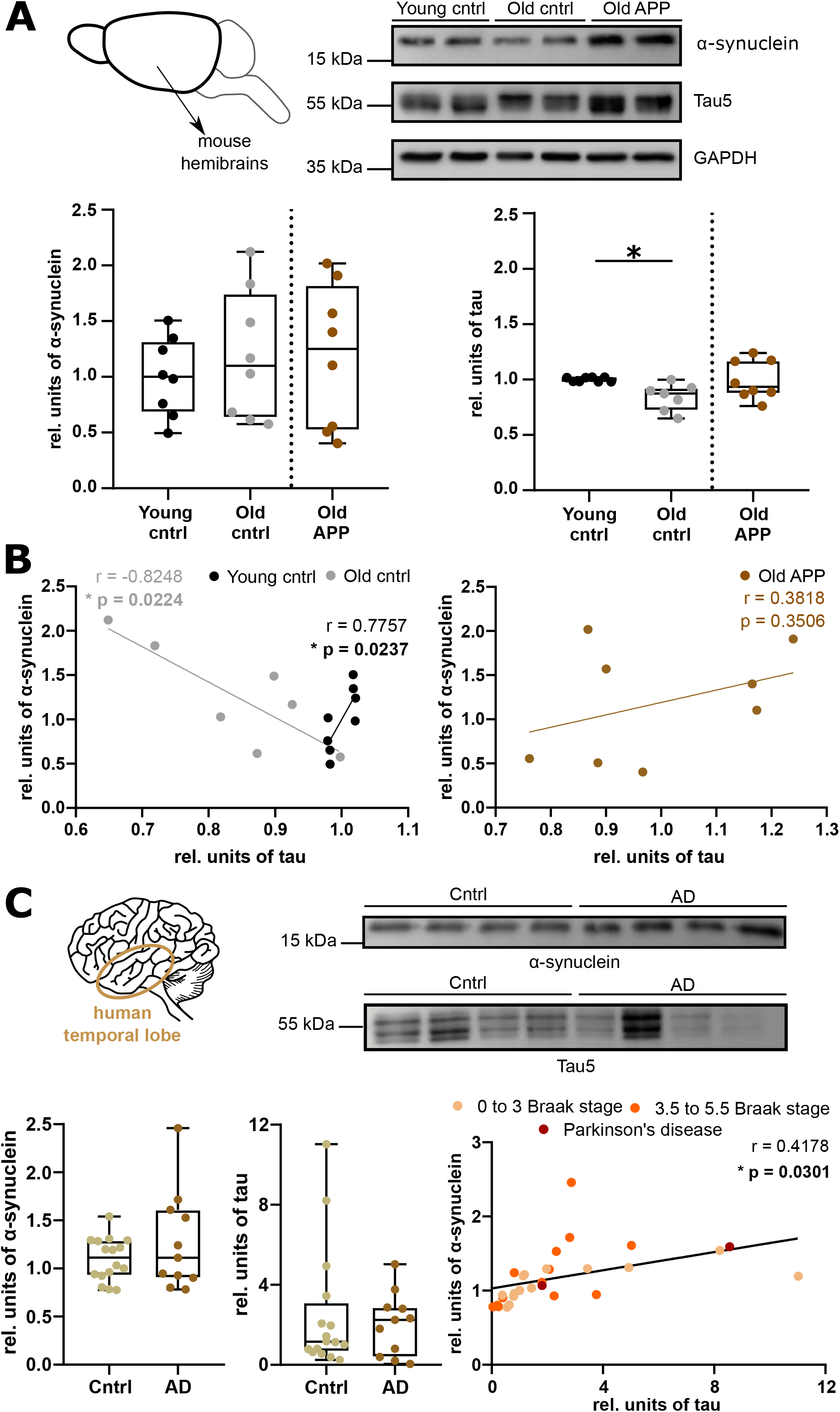
Tau and α-syn reveal an inverse regulation in old mice under physiological conditions, which is disrupted during brain pathology. **A**. Pictogram representing a mouse hemibrain, followed by representative western blot bands of the expression of α-syn, tau and GAPDH, which was used to normalize protein load in young (3-4-months-old) control and old (20-24-months-old) controls, and APP mice. Molecular mass standards are indicated. Below, box plots displaying the quantification of α-syn and Tau expression. **B**. Scatter plot on the left correlates α-syn and Tau expression from young and old control mice. On the right, correlation of α-syn and Tau expression from old APP mice. n = 8 young control, n = 8 old control, and n = 8 old APP hemibrains used for α-syn expression measurements; n = 8 young control, n = 7 old control, and n = 8 old APP hemibrains used for Tau expression measurements and expression correlation. Statistical analysis employed unpaired two-tailed t-test with Welch’s correction and Pearson’s correlation. *p<0.05. **C**. Schematic representation of a human brain indicating that temporal lobe samples were employed. On the right, representative western blot bands of α-syn and Tau expression in control and Alzheimer’s disease (AD) patients. Below, box plots displaying the relative amount of α-syn and Tau expression after normalization to the protein load of each sample. On the right, scatter plot showing the correlation between the expression of the proteins in control, AD, and Parkinson’s disease (PD) post-mortem samples. n = 16 control, n = 11 AD, and n = 2 PD human temporal lobe lysates. Statistical analysis employed unpaired two-tailed t-test and Pearson’s correlation was used, excluding PD samples. *p<0.05.

### Tau and α-syn levels are uncorrelated in mice with amyloidosis and positively correlated in the elderly human brain

Most patients with Parkinson’s and Alzheimer’s have mixed types of protein deposits containing tau and α-syn in the brain. To determine whether a negative correlation between tau and α-syn levels also exists under disease conditions, we investigated a possible correlation between the levels of the two proteins in a mouse model of amyloidosis and AD patients. Interestingly, when examining lysates from hemibrains of old mice with chronic amyloidosis (Hrynchak et al., 2020), we observed no decrease in tau protein with age and the negative correlation between tau and α-syn was also absent (Figs. 5A, B). We extended our analysis to include brain samples from mostly elderly human subjects. There was no statistical difference in relative tau or α-syn levels between these groups of human samples clinically confirmed as AD patients and controls. However, when we correlated the two protein levels for each sample, we observed a significant positive correlation. We also obtained two human samples from PD patients. Due to the small number of samples, we did not include them in the statistics, but we found that the samples with synucleinopathy also matched the linear regression obtained from the other samples. Thus, the data may suggest that a decrease in tau levels and compensation by α-syn is beneficial during physiological aging and may stop functioning in diseased brain states.

## Discussion

Each phase of nervous system development requires the presence of sophisticated and tightly regulated molecular machinery to achieve and maintain a structure that provides adequate behavioral performance. This also applies to the maturation of the dendritic processes because the dendritic architecture fundamentally determines the functional capacity of neurons. During early postnatal development significant dendritic process formation takes place in various brain areas (Kolb and Fantie, 1997) (Richards et al., 2020) and the formation and pruning of dendritic processes determine the density of the neural network in which individual neurons become embedded. Given the unique function and shape of the dendritic arbor, the integrity of the microtubule cytoskeleton and proper transport processes are crucial during dendritic process formation (Bramham and Wells, 2007; Penazzi et al., 2016).

The present findings (1) identify a more complex branching pattern of the maturing dendritic tree of hippocampal pyramidal cells in the tissue of Tau KO mice. They indicate that (2) the highly elaborate dendritic tree likely evolved due to a more directional transport mechanism that we recognized after KIF1A mediated transport. We show that (3) the transport pathways, the microtubule lattice, are longer and straighter in Tau KO mice. Our results suggest that (4) α-syn is an accomplice in modulation of transport properties, as we found highly increased levels of α-syn in Tau KO hippocampi and that overexpression of α-syn in control hippocampal tissue mimics the transport properties observed in cells lacking tau protein. Finally, we found that (5) α-syn and tau are negatively correlated during physiological aging, but not under disease conditions.

Microtubules and motor proteins that move along microtubule tracks play an essential role to accurately control the intracellular transport of various cellular cargo. Microtubules are also the core cytoskeletal elements that are determining dendritic structure by exerting force in order to create, stabilize, and remodel dendrites during development (Janke and Magiera, 2020; Kapitein and Hoogenraad, 2015; Poulain and Sobel, 2010). Tau is a well-known microtubule-associated protein. *In vitro* it had been shown that tau protein promotes microtubule nucleation and elongation from tubulin which would not assemble in the absence of MAPs (Cleveland et al., 1977; Witman et al., 1976). Tau also modulates the dynamic instability of tubulin assembly by affecting the rates of polymerization, transition into catastrophic depolymerization, and depolymerization (Drechsel et al., 1992; Trinczek et al., 1995). Sense and Antisense transfection into PC12 cells showed that tau influences net microtubule assembly, neurite outgrowth and stability (Esmaeli-Azad et al., 1994). Previously, we showed that acute overexpression of human tau in PC12 cells led to increased levels of microtubules in the polymer form (Janning et al., 2014). However, here we show that live-imaging of cultured hippocampal cells from controls and cells lacking tau showed no significant difference in dendritic microtubule dynamics. Presence of tau has been shown within dendrites and at the post-synapse, both under physiological and pathological conditions (Ittner et al., 2010). However, in white matter three times more tau was identified compared to the grey matter, indicating a much higher abundance of tau in the axonal processes versus the somatodendritic compartment (Binder et al., 1985). However, later it was shown that the antibody recognized a dephosphorylated form of tau and total tau was more abundant in dendrites than originally thought (Papasozomenos and Binder, 1987). The major interaction partners of tau, the microtubules are also showing differences with respect to orientation, density and dynamics, which is presumably favoring modulation by this lower tau amounts. Therefore, in the somatodendritic compartment of neuronal cells where tau is knocked out, microtubule dynamics and structure are affected by a different molecular environment. Using non-biased mass spectrometry analysis, we found no evidence for an increase of microtubule-related structural proteins for compensatory mechanisms. However, we found that some proteins from both kinesin and dynein family were elevated in the hippocampal tissue of Tau KO mice. Motor proteins use the microtubule cytoskeleton as a road for transport of vesicles and organelles (Barlan and Gelfand, 2017) and the transport relies on the state of dynamics of microtubules (Masucci et al., 2022; Yang et al., 2017). Motor proteins themselves could influence microtubule structure and dynamics (Yogev et al., 2017). Microtubule organization, in turn, determines the progression of axonal transport: cargoes pause at polymer termini, suggesting that switching microtubule tracks is rate limiting for efficient transport. Cargo run length is determined by the length of microtubules (Yogev et al., 2016). Therefore, it is also important to understand potential structural alterations of microtubules not just their dynamics.

For this study, we have overcome a major technical barrier to analyze structural parameters of microtubules in neuronal cells. Previous attempts relied on the reconstruction of microtubule length from serial sections of electron microscopy images, primarily from simple organisms, such as *C. elegans* (Chalfie and Thomson, 1979). Information on the length of microtubules from cultured hippocampal cells, including the dendritic domain, later emerged by serial reconstruction of the microtubule array from electron micrographs (Yu and Baas, 1994) or by following the growth of microtubules after microinjection of biotin-labeled tubulin (Wang et al., 1996). These studies report an average microtubule length of just over 3 µm and the longest microtubules (with very low abundance) reaching over 10 µm. We found similar values for the mean microtubule length in both conditions analyzed using super resolution light microscopy based on 3D DNA-PAINT. However, total microtubule length was significantly increased in cells lacking tau protein. We could also observe that microtubules were straighter and more parallel arranged suggested by the lower number of junctions observed between microtubule cylinders situated close to each other. All this allows motor proteins to run longer towards the designated direction.

Although the ratio between minus-and plus-end out microtubule orientations changes depending on the neuron type and even differs between regions of the same dendrite (van de Willige et al., 2016), vertebrate neuronal cells favor microtubules with mixed polarity in dendrites in more or less equal proportions (Stone et al., 2008). Therefore, even motor proteins that have a dedicated orienting motion to a particular microtubule end would show mixed motions in these processes. We observed motion of KIF1A in dendritic processes from and towards the neuronal soma. In tissue cells lacking tau protein however, the processivity of the movement was increased. Individual tracks revealed that this increased processivity is due to less changes in the direction of the motor protein, granting a more forward movement.

We have encountered high overexpression of α-syn in Tau KO hippocampal tissue by mass spectrometry and western blot analysis. Most of the previous publications suggested that α-syn plays a role in various pre-synaptic processes because it is enriched in the presynaptic terminals throughout the adult mammalian brain (Petersen et al., 1999; Twohig and Nielsen, 2019). However, several studies demonstrated interaction between physiological α-syn or its disease-relevant, aggregated conformers and microtubule elements (Alim et al., 2002; Cartelli et al., 2016; Seebauer et al., 2022). In agreement, we observed that α-syn is decorating dendritic microtubules independent of the presence of tau. A potential effect on microtubule structure could explain that overexpression of α-syn in control tissue cells resembled transport properties seen in cells without tau protein.

Both tau and α-syn are thought to be intrinsically disordered proteins (Brandt et al., 2020; Zhang et al., 2020), making them promiscuous with respect to interaction partners and allowing for a broader determination of their function. We demonstrate a previously unappreciated role for dendritic tau and α-syn in shaping transport in maturing dendritic processes in the hippocampus. We also show that during young age tau and α-syn are positively correlating with each other in mice. However, physiological aging in mice induces a negative correlation between tau and α-syn, which is lost when mice exhibit disease-relevant conditions, suggesting a drive towards a condition that is providing a molecular milieu for neurons during dendritic maturation.

## Figure legends

**Supplementary Figure 1. GO-term representation of major functions for differentially regulated proteins in Tau KO hippocampi.** GO-terms significantly enriched (p<0.01) for lists with up-or down-regulated proteins in Tau KO are shown in purple or grey color, respectively. GO-terms significantly enriched (p<0.01) for lists with both up-and down-regulated proteins in Tau KO are shown in yellow. The size of the nodes illustrates statistical significance, the larger nodes correspond to higher significance of GO-term enrichment. The fractions of proteins are shown inside for upregulated Tau KO dataset, up-regulated proteins in Tau KO hippocampi are shown in purple and down-regulated – in grey. The network representation was created in ClueGO plugin for Cytoscape.

**Supplementary Figure 2. α-synuclein distribution in hippocampal neurons at 28 DIV.** Exemplary micrographs showing the z-projection of a control and a Tau KO hippocampal primary neuron immunostained with anti-α-synuclein antibody at 28 DIV after NP-40 extraction (a,b). A z-plane displaying the inner part of the dendrite was selected (c,d). Selected z-planes were zoomed in to show the distribution of α-synuclein decorating microtubule bundles (red arrows) and presynaptic buttons (yellow arrows) (e,f).

## Acknowledgement

This work was supported by Deutsche Forschungsgemeinschaft (SFB 944, projects Z, PI 405/10-1, the DFG Facility iBiOs & INST 190/182-1 to RK) and (DFG BR1192/14-1 to RB).

