## Supplemental material for "Tau and α-synuclein shape microtubule organization and microtubule-dependent transport in neuronal dendrites"

Table 1. List of the human samples that were used for the study

| case number | sex | age (yeasrs) | Braak stage | postmortem (h) | cause of death |
|---|---|---|---|---|---|
| AD1 | w | 86 | 3,5 | 120 | cardiovascular failure |
| AD2 | m | 93 | 4,0 | 49 | lung cancer |
| AD3 | m | 78 | 4,0 | 51 | pneumonia |
| AD4 | w | 75 | 3,5 | 105 | cardiovascular failure |
| AD5 | w | 77 | 4,0 | 62 | cardiovascular failure |
| AD6 | w | 79 | 4,5 | 19 | pulmonary embolism |
| AD7 | m | 73 | 3,5 | 105 | cardiovascular failure |
| AD8 | w | 84 | 5,5 | 72 | pneumonia |
| AD9 | w | 77 | 5,5 | 34 | cardiovascular failure |
| AD10 | w | 92 | 5,5 | 48 | pneumonia |
| AD11 | w | 82 | 5,5 | 48 | pneumonia |
| CO1 | w | 74 | 2 | 33 | cardiovascular failure |
| CO2 | w | 67 | 2,5 | 54 | cardiovascular failure |
| CO3 | w | 83 | 4 | 48 | cardiovascular failure |
| CO4 | m | 73 | 0,5 | 9 | cardiovascular failure |
| CO5 | w | 76 | 2,5 | 120 | cardiovascular failure |
| CO6 | w | 85 | 3,5 | 57 | suicide |
| CO7 | w | 79 | n.a. | 33 | multiorgan failure |
| CO8 | m | 73 | n.a. | 24 | cardiovascular failure |
| CO9 | m | 84 | n.a. | 36 | cardiovascular failure |
| CO10 | m | 76 | 0 | 46 | cardiovascular failure |
| CO11 | m | 60 | n.a. | 38 | cardiovascular failure |
| CO12 | m | 51 | n.a. | 48 | pulmonary embolism |
| CO13 | m | 40 | n.a. | 35 | kidney failure |
| CO14 | f | 38 | n.a. | 38 | cardiovascular failure |
| CO15 | m | 31 | n.a. | 51 | cardiovascular failure |
| CO16 | m | 25 | n.a. | 76 | liver failure |

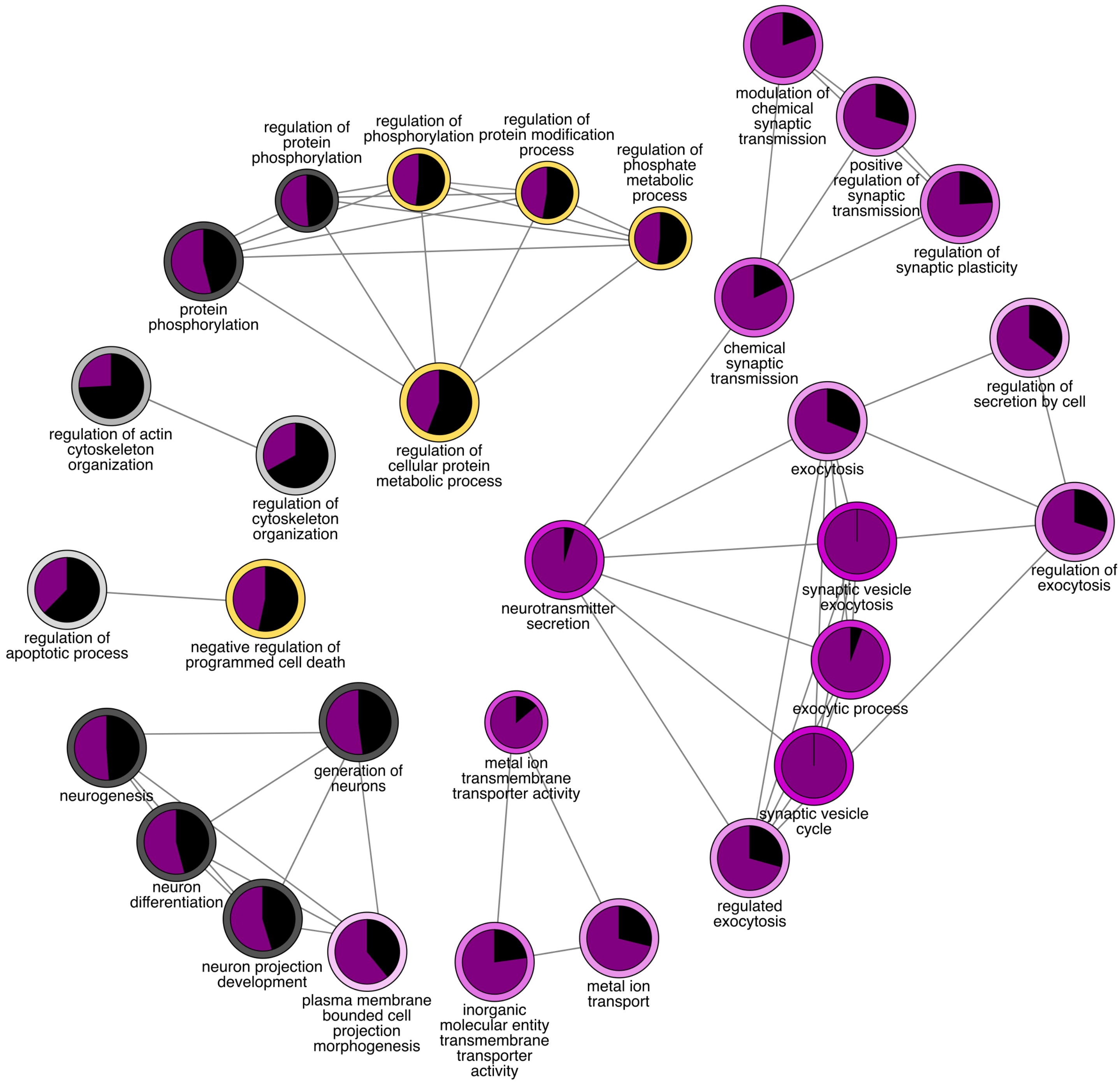


**Supplementary Figure 1. GO-term representation of major functions for differentially regulated proteins in Tau KO hippocampi.**

GO-terms significantly enriched (p<0.01) for lists with up- or down-regulated proteins in Tau KO are shown in purple or grey color, respectively. GO-terms significantly enriched (p<0.01) for lists with both up- and down-regulated proteins in Tau KO are shown in yellow. The size of the nodes illustrates statistical significance, the larger nodes correspond to higher significance of GO-term enrichment. The fractions of proteins are shown inside for upregulated Tau KO dataset, up-regulated proteins in Tau KO hippocampi are shown in purple and down-regulated – in grey. The network representation was created in ClueGO plugin for Cytoscape.

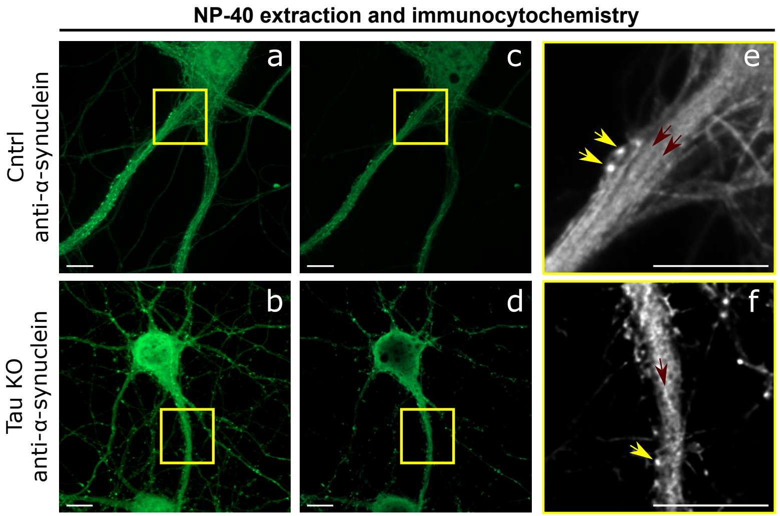


**Supplementary Figure 2. α-synuclein distribution in hippocampal neurons at 28 DIV.**
Exemplary micrographs showing the z-projection of a control and a Tau KO hippocampal primary neuron immunostained with anti-α-synuclein antibody at 28 DIV after NP-40 extraction (a,b) . A z-plane displaying the inner part of the dendrite was selected (c,d). Selected z-planes were zoomed in to show the distribution of α-synuclein decorating microtubule bundles (red arrows) and presynaptic buttons (yellow arrows) (e,f).
